# Intracellular level of *S. cerevisiae* Rad51 is regulated via proteolysis in a SUMO- and ubiquitin-dependent manner

**DOI:** 10.1101/2022.05.31.494133

**Authors:** Justyna Antoniuk-Majchrzak, Anna Długajczyk, Tuguldur Enkhbaatar, Joanna Kaminska, Marek Skoneczny, Daniel J. Klionsky, Adrianna Skoneczna

**Author notes:** Adrianna Skoneczna. Institute of Biochemistry and Biophysics, Polish Academy of Sciences, Pawińskiego 5A, 02-106 Warsaw, Poland, **Email:**.

## Abstract

Among various DNA lesions, the DNA double-strand breaks are particularly deleterious; especially, when an error-free repair pathway is unavailable, and the cell takes the risk of using the error-prone recombination pathways to repair the DNA breaks, resume the cell cycle, and continue growth. The latter comes at the expense of decreased well-being of the cells due to genome rearrangements. One of the major players involved in recombinational repair of DNA damage is Rad51 recombinase, a protein responsible for presynaptic complex formation. We previously noticed that the level of this protein is strongly increased when illegitimate recombination is engaged in repair. The regulation of Rad51 protein turnover is not known; therefore, we decided to look closer at this issue because we expect that an excessively high level of Rad51 may lead to genome instability. Here we show that the level of Rad51 is regulated via the ubiquitin-dependent proteolytic pathway. The ubiquitination of Rad51 depends on multiple E3 enzymes, including SUMO-targeted ubiquitin ligases. We also demonstrate that Rad51 can be modified by both ubiquitin and SUMO. Moreover, these modifications may lead to opposite effects. Ubiquitin-dependent degradation depends on Rad6, Rad18, Slx8, Dia2 and the anaphase-promoting complex. Rsp5-dependent ubiquitination leads to Rad51 stabilization.

## Introduction

The maintenance of genetic information is essential for cells but not easy to achieve due to constant environmental and metabolic threats. Particularly deleterious for genome integrity are stresses causing double-strand breaks in DNA because this type of damage may lead to DNA rearrangements or loss of parts of chromosomes. To avoid such scenarios and restore vital genetic information, the repair pathways evolved, reassuring the re-connection of the broken DNA strands. In the yeast *Saccharomyces cerevisiae*, the most accurate repair pathway dedicated to double-strand break repair is homologous recombination. One of the proteins essential for this repair is the Rad51 recombinase. Rad51 belongs to the RecA family of recombinases (1). Due to its capacity to bind single- and double-stranded DNA, DNA-dependent ATPase activity, ability to form a filament on DNA, and feedback interactions with various proteins engaged in DNA repair (Rad54, Rad52, Sgs1, replication protein A/RPA complex, etc.), the recombinase Rad51 executes the critical early step of homologous recombination: the search for homologous DNA to serve as a template during the repair of DNA double-strand breaks (2-6). The *rad51* mutants display replication defects and chromosomal instability (7). The orthologs of yeast *RAD51* gene have been identified in various organisms, including humans (8). In vertebrates, the absence of Rad51 leads to embryonic lethality (9). Moreover, mutations in human RAD51 are linked to breast cancer (10,11) and Fanconi anemia (complementation group R, FANCR) (12,13).

Rad51 is involved in mitotic and meiotic recombination; however, its role is slightly different in each process. Because Rad51’s homolog Dmc1 is present during meiosis, the proteins share their responsibilities. While Dmc1 is responsible for interhomolog recombination, Rad51 promotes Dmc1 presynaptic filament assembly and participates in intersister repair, leading to non-crossover products (14-16).

Rad51 activity is regulated on several levels. Besides interactions with DNA and various DNA repair proteins, Rad51 activity depends on post-translational modifications, e.g., phosphorylation. The Cdc28-dependent phosphorylation of Ser125 and Ser375 at the G2/M border of the cell cycle promotes the DNA binding affinity of Rad51 (17). In turn, Mec1-dependent phosphorylation on Ser192 residue is required for two functions of Rad51: DNA-binding and ATPase activity (18). The abundance of Rad51 in the cell also influences its function. The availability of Rad51 is determined by the level of expression of the *RAD51* gene, which rises in a cell cycle-dependent manner during G1/S transition and after exposure to genotoxic stress when demand for Rad51 protein increases (19, 20). But the abundance of Rad51 in the cell strongly depends on its half-life life, which should be tightly controlled to regulate and resume the homologous recombination repair pathway. Indeed, the high level of Rad51 during replication stress leads to an increased usage of illegitimate recombination causing frequent genome rearrangement (21).

Here we showed that Rad51 protein level is controlled by proteolytic digestion. Its half-life changes in response to environmental conditions, e.g., upon genotoxic stress. We also identified several ubiquitin-conjugating enzymes (E2), ubiquitin ligases (E3) and SUMO (E3) ligases affecting the Rad51 cellular level. We showed that Rad51 is post-translationally modified by ubiquitin and SUMO. Interestingly, different patterns of Rad51 ubiquitination/SUMOylation were visible in various conditions. Results showed that a proteolytic pathway controls the Rad51 level in the cell in a ubiquitin- and SUMO-dependent manner. A range of factors is involved in the regulation of this process, signifying the need for precise Rad51 regulation.

## Results

### Rad51 protein level in the cell is actively regulated via ubiquitin-dependent proteolysis

One of the ways to regulate protein functioning in the cell is to control its abundance. The fastest way to do it is to limit its proteolysis when the protein is needed. We asked if the half-life of Rad51 is changed under genotoxic stress conditions when there is a higher demand for its activity. Accordingly, we analyzed the stability of Rad51, following the addition of cycloheximide (CHX) to inhibit protein translation. We found that Rad51 degradation displayed a steady rate of turnover in the wild-type strain following treatment with CHX (Figure 1 A, B). When these cells were additionally treated with methyl methanesulfonate (MMS) or zeocin to induce DNA damage the degradation of Rad51 was reduced. The results suggested that Rad51 degradation is actively regulated in response to DNA damage.

**Figure 1.**
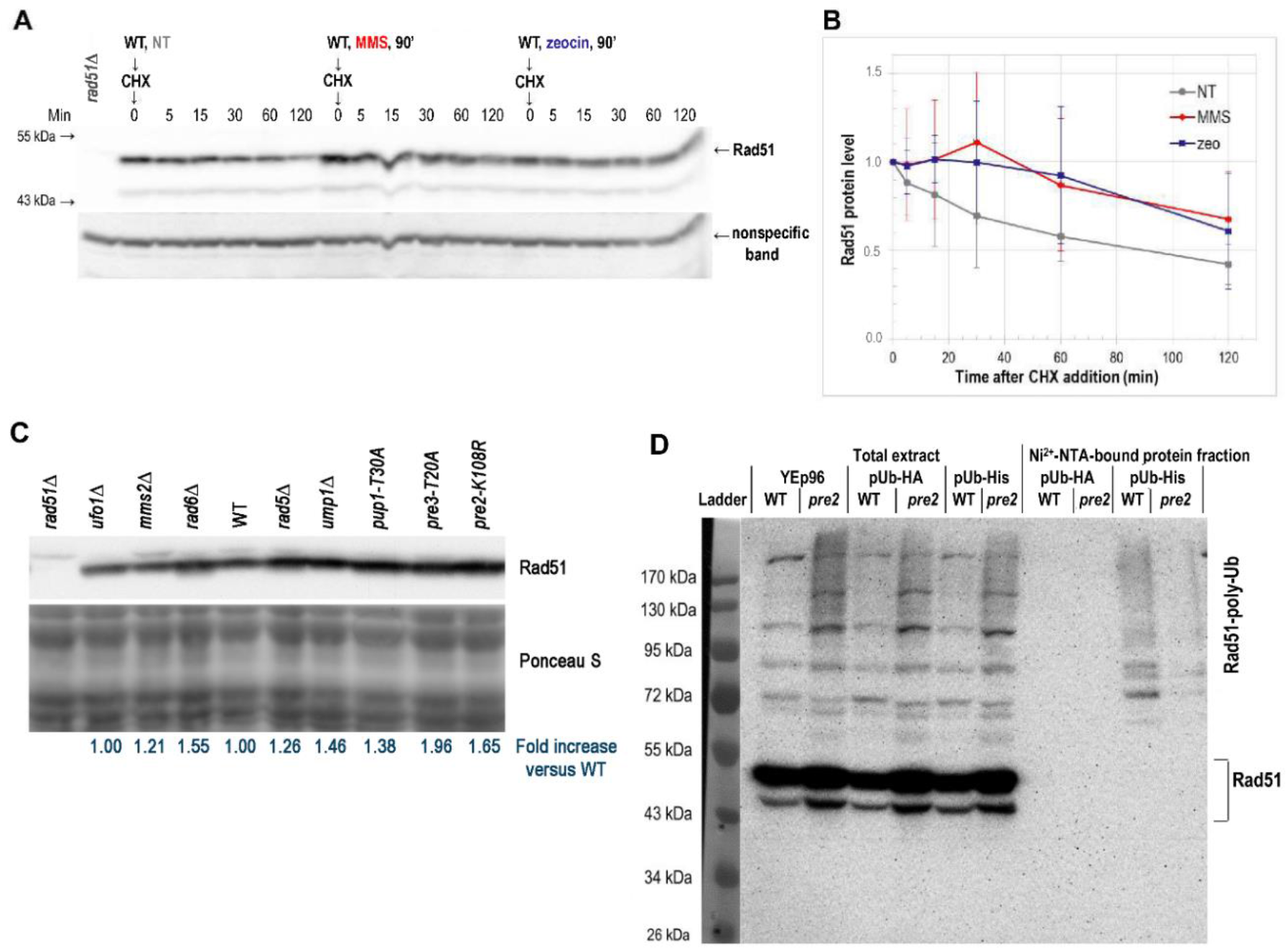
Rad51 level is regulated by ubiquitin-dependent proteolysis. (**A**) Rad51 stability is changed after genotoxic stress. A culture of the WT strain (BY4741) was divided into three parts: control (NT), treated with 0.03% MMS (MMS), and treated with 50 mg/ml zeocin (zeo). After 90 min of incubation, cycloheximide (CHX) was added to a final concentration of 0.5 mg/ml. Protein samples were harvested at the indicated time points, and immunoblotting was performed (**A**). The immunoblotting results shown in (**A**) were quantified and plotted in (**B**). Data from three independent biological repetitions. (**C**) The steady-state level of Rad51 increases in proteasomal mutants. Exponentially growing cells of the WT (WCG4a), *pre2-K108R* (YWH24), *pre3-T20A* (YUS1), *pup1-T30A* (YUS4), *ump1Δ* (YAS13), *rad5Δ* (YJM26), *rad6Δ* (YAZ10), *ufo1Δ* (YJD1), and *mms2Δ* (YJM56) strains were assayed by western blot to monitor the Rad51 level. The Rad51 signal was normalized to the proteins visualized by Ponceau S staining. The average fold increase versus the WT Rad51 level, which was set to 1.00, was then calculated (blue numbers below the blot). (**D**) Analysis of Rad51 ubiquitination *in vivo* by His-ubiquitin affinity-isolation assay. WT (YJK1) and *pre2-K108R* (YJK2) strains bearing an empty vector YEp96 or a vector encoding His6-ubiquitin and HA-ubiquitin (control) from plasmids YEp96-Ubi-His (pUb-His) (22) and YEp112 (pUb-HA) (23), respectively were grown in the presence of Cu^2+^ ions and collected. Ubiquitinated proteins were purified on Ni^2+^-NTA sepharose from total extracts and analyzed by western blot using anti-Rad51 antibodies. The position of Rad51 and poly-ubiquitinated Rad51 (Rad51-poly-Ub) were indicated on the right. Molecular mass markers were displayed on the left.

Cells wield multiple proteolytic systems to carry out protein degradation. One of the most common pathways used for this purpose is ubiquitin-dependent proteolysis via the 26S proteasome. To answer the question of whether Rad51 is a substrate for the proteasome, we measured the steady-state level of Rad51 in yeast mutants defective in various proteasomal activities. We exploited the *pre2-K108R, pre3-T20A*, and *pup1-T30A* mutant strains, where chymotrypsin-like, trypsin-like, and peptidyl-glutamyl peptide-hydrolyzing (PHGH) activities of 20S proteasome core were inactivated, respectively (22). In proteasome mutant strains there was an increased level of Rad51 (Figure 1 C). A similar effect was visible in *ump1Δ* cells lacking proteasome maturase (23). We therefore concluded that Rad51 is degraded by the proteasome.

The proteins that undergo proteasomal degradation first have to be tagged with ubiquitin. Thus, we took advantage of the His-tagged ubiquitin system to check whether Rad51 is modified by ubiquitin. Using Ni^2+^-NTA sepharose, we purified ubiquitinated proteins from the strains carrying a plasmid expressing a His-tagged ubiquitin variant under the control of the inducible *CUP1* promoter (24). The Ni^2+^-NTA-bound fraction of proteins, i.e., proteins modified with ubiquitin, and total cell extract, were then separated by SDS-PAGE and analyzed by western blot. Anti-Rad51 antibodies detected poly-ubiquitinated variants of Rad51 among the Ni^2+^-NTA-bound protein fraction (Figure 1 D) from cells expressing His-tagged ubiquitin. In contrast, we did not detect any poly-ubiquitinated Rad51 in the strain expressing HA-tagged ubiquitin as a negative control. Thus, the Rad51 protein is poly-ubiquitinated.

We performed a similar analysis with the *pre2-K108R* strain, expecting better visibility of Rad51 ubiquitinated forms due to decreased turnover; however, we detected a reduced level of the poly-ubiquitinated species (Figure 1 D). Based on the total cell extract it was clear that there was a relative accumulation of Rad51, including the poly-ubiquitinated forms, in *pre2-K108R* cells. One possible explanation would be that the *pre2-K108R* cells accumulate ubiquitinated proteins, due to defects in proteasome-dependent degradation, and these substrates compete for binding to the Ni^2+^-NTA sepharose. Therefore, although the number of Rad51 poly-ubiquitinated forms increases in the *pre2-K108R* mutant, they cannot be visualized efficiently in such an analysis. In agreement with this hypothesis would be our observation that the flow-through fraction still contains a substantial level of higher molecular weight forms of Rad51 (data not shown). Interestingly, the pattern of poly-ubiquitinated forms of Rad51 in total extracts and Ni^2+^-NTA-bound protein fraction is slightly different, which suggests His-tagged ubiquitin does not necessarily substitute for the endogenous ubiquitin perfectly in all places open to this kind of modification or that the His-tagged ubiquitin does not form poly-ubiquitin chains with the same efficiency as native ubiquitin. Another explanation for the lower level of Rad51 poly-ubiquitinated forms visible in the Ni^2+^-NTA-bound fraction in the *pre2-K108R*-derived sample would be that there is a compensatory effect; when the Rad51 level is too high, cells start to use another system enabling its degradation. Having a backup system for Rad51 degradation would be of biological importance because a high level of Rad51 leads to an increased usage of illegitimate recombination, causing genome instability and elevated mortality of the cells (21).

### Multiple E2 and E3 enzymes participate in the in vivo degradation of Rad51

Because Rad51 is poly-ubiquitinated, and our preliminary results showing increased levels of Rad51 in the strains lacking enzymes involved in protein ubiquitination were promising (Figure 1 C), we asked which E2 and E3 enzymes are responsible for that modification. To answer this question, we first performed in silico studies. We searched the *Saccharomyces* Genome Database (SGD) (RRID:SCR_004694) looking for proteins responsible for proteolysis or contributing to its regulation. In addition, we explored the *RAD51* dataset deposited in the Biological General Repository for Interaction Datasets (BioGRID) (RRID:SCR_007393), looking for Rad51 interactors. Then, we compared the two obtained datasets; overlapping candidates were examined further. We examined the effect of null or conditional mutants in the BY4741 background on the level of Rad51 (Figure 2). The screen confirmed the dependence of Rad51 levels on proteasomal activities because a twofold increase of Rad51 was observed in all assayed *pre2* mutants and the *ump1Δ* strain, in agreement with our initial findings (Figure 1 C). The screen further implicated a group of E2 and E3 enzymes that were likely to be involved in posttranslational Rad51 modifications because the lack of their proper function led to an increased Rad51 steady-state level. In particular, these included strains with mutations in Dia2 and Cdc4, the subunits of respective Skp1-Cullin1-F-box protein (SCF) ubiquitin ligase complexes; RING finger proteins Rad5 and Rad18, as well as the SUMO-targeted ubiquitin ligase (STUbL) Slx8 (Figure 2 A). Similar phenotypes were seen in strains lacking: (i) ubiquitin-conjugating enzymes Rad6, Mms2, and Ubc13 (cooperating with Rad18 or Rad5); (ii) subunits of the SCF complex Skp1, Cdc34, and Cdc53; (iii) or SUMO-ligases, such asWss1 or strain lacking both, Siz1 and Nfi1/Siz2 SUMO-ligases (Figure 2 A, Figure 2 B). Moreover, these results pointed out the importance of an anaphase-promoting complex (APC) for Rad51 stability because in *apc11-22* and *cdc23-1* strains, carrying mutations in APC subunits, the level of Rad51 was also elevated. Mutants that showed the significant decrease in the Rad51 level were strains lacking a functional Rsp5 protein (a NEDD4 family E3 ubiquitin ligase) and the Prp19 protein whose U-box domain possesses E3 ubiquitin ligase activity (Figure 2 A).

**Figure 2.**
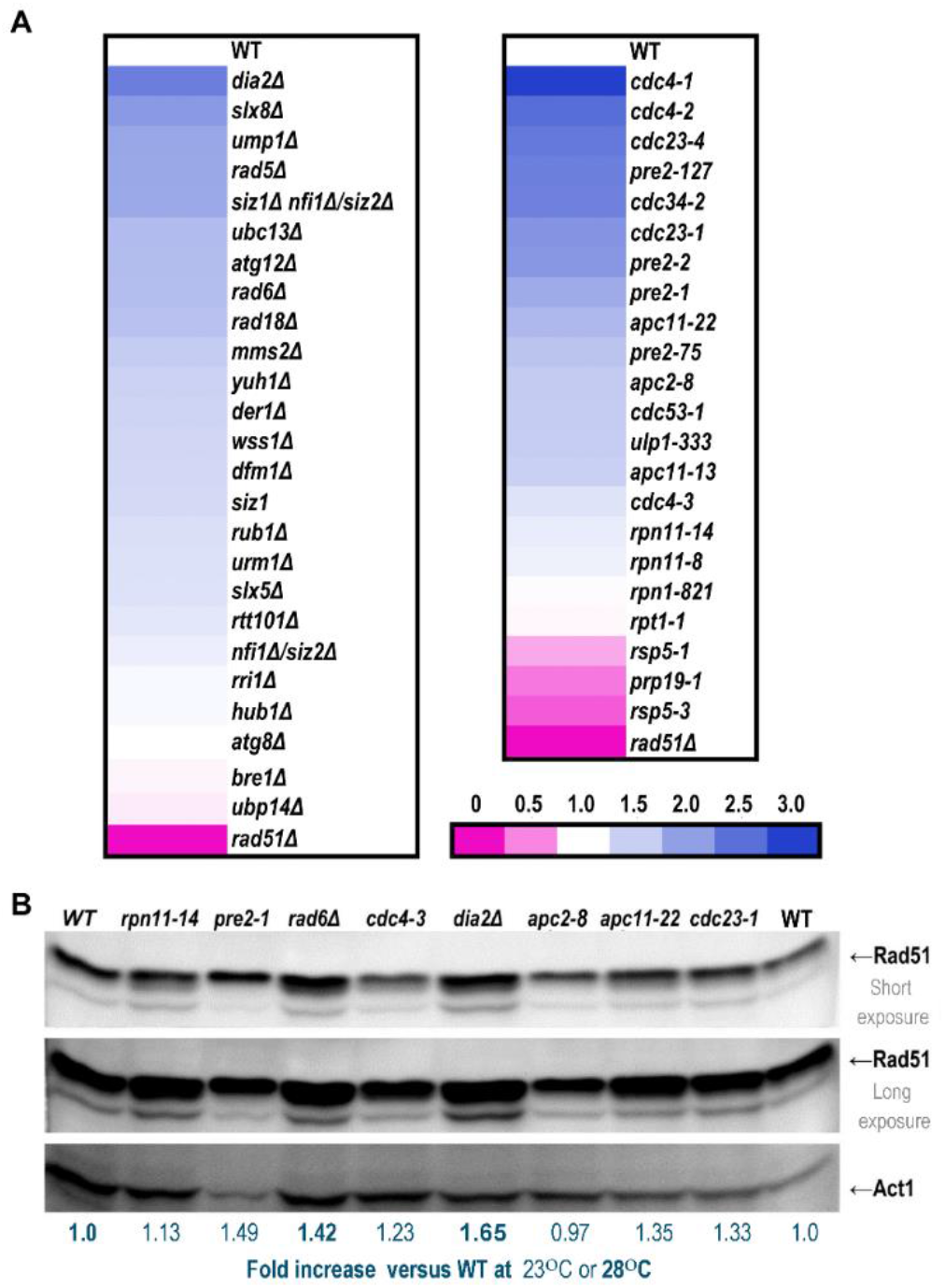
Multiple E2 and E3 enzymes influence the Rad51 cellular level. (**A**) Heat map showing the fold change of the Rad51 level in the indicated mutants relative to the level of this protein in the wild-type control. Strains in the BY4741 background, bearing defects in the genes encoding proteins involved in protein stability control via the ubiquitin-dependent degradation pathway, were assayed by western blot. Protein extracts were prepared from cells in the exponentially growing phase. From 2 to 6 independent biological replicates were made. The Rad51 signal was normalized to actin, a nonspecific band, or Ponceau S-stained proteins. The average fold change of Rad51 level relative to the WT cells was shown. The left and right panels show results obtained at 28°C or 23°C, respectively. (**B**) Western blot showing the dependence of Rad51 degradation on selected enzymes from the ubiquitin-dependent degradation pathway. Deletion strains *rad6Δ* and *dia2Δ* and the WT strain shown on the left were grown 28°C, and strains carrying point mutations in the essential genes *APC2, APC11, CDC23, CDC4, PRE2*, and *RPN11* and the WT strain shown on the right were grown at 23°C. Act1 is shown as a loading control. Blue numbers below the blot represent for the presented western blot result the fold increase versus the WT Rad51 level, which was set to 1.00. For the data obtained at 23°C regular fonts were used, while for the data obtained at 28°C bold fonts were used.

### SUMOylation contributes to Rad51 protein level regulation

As shown in Figure 1 D, the Rad51 protein is poly-ubiquitinated *in vivo*. However, the mono-ubiquitinated form of Rad51 was not visible on the western blots among Ni^2+^-NTA-bound protein fractions. While studying the Rad51 western blots signals, we noticed another characteristic feature of this protein: it is represented by multiple bands. The weak band migrated just as expected for Rad51 molecular mass (about 43 kDa), and the more pronounced forms migrated around 50 kDa. For this mobility shift, the post-translational modification of Rad51 might be responsible. According to published data, several amino acid residues within the Rad51 sequence are subject to phosphorylation. Because the shift in Rad51 mobility visible on western blots can be estimated as corresponding to a 7- to 8-kDa change in molecular mass of this protein, the probability that phosphorylation status is responsible seems unlikely. In contrast, the mass difference suggests a ubiquitin-like modifier. To test such a hypothesis, we used a series of yeast mutants lacking genes encoding various ubiquitin-like modifiers: Atg12 (involved in autophagy (25)), Hub1 (involved in budding and mating processes (26, 27)), Rub1 (involved in neddylation of various substrates, e.g., cullins (28)), and Urm1 (involved in glucose limitation response and oxidative stress response (29)). However, both the level of Rad51 and the position of the protein following SDS-PAGE were similar in all analyzed strains (Figure 3 A).

**Figure 3.**
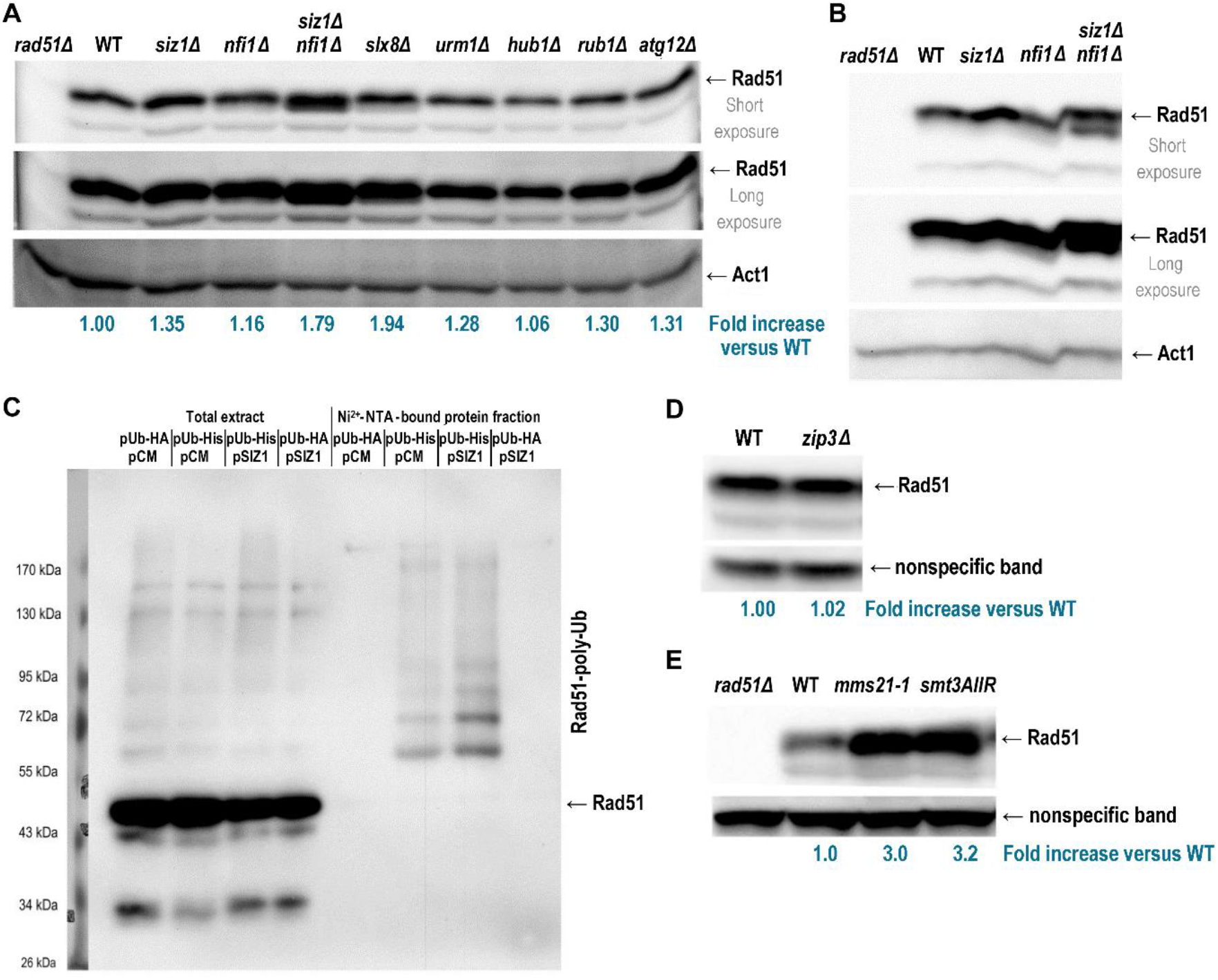
SUMOylation promotes Rad51 destabilization. (**A**) Rad51 accumulates in mutants lacking SUMO E3 ligases or STUbL complex subunit Slx8. Strains in the BY4741 background lacking the indicated genes were cultivated to an exponential growth phase, then were subjected to western blot analysis to detect Rad51. At least three independent biological replicates were made. The average fold increase versus the WT Rad51 level, which was set to 1.00, was then calculated (blue numbers below the blot). (**B**) A strong, additional Rad51-specific band was observed on the western blot in the *siz1Δ nfi1/siz2Δ*-derived samples. (**C**) Analysis of Rad51 ubiquitination *in vivo* by His-ubiquitin affinity-isolation assay. A WT (YJK1) strain bearing the plasmid YEp96-Ubi-His (pUb-His) (22) or YEp112 (pUb-HA) (23) and pCM188-SIZ1 (overexpressed Siz1) or the pCM188 empty vector, respectively, were grown in the presence of Cu^2+^ ions and collected. Ubiquitinated proteins were purified on a Ni^2+^-NTA Sepharose column from total extracts and analyzed by western blot using anti-Rad51 antibodies. Rad51 and poly-ubiquitinated Rad51 (Rad51-poly-Ub) were indicated on the right. Molecular mass markers were shown on the left. (**D**) A *zip3Δ* strain displays a Rad51 level similar to the WT strain. The WT and *zip3Δ* strain-derived samples were prepared and probed as in (**A**). (**E**) Rad51 accumulated in cells lacking Mms21 SUMO-ligase and in cells bearing an *smt3AllR*-encoding SUMO variant deprived of all its lysine residues, therefore, disabling chain structure formation. The WT, *mms21-1*, and *smt3AllR* strain-derived samples were prepared and probed as in (**A**). Act1 serves as a loading control in (A), (B) and (E), and a nonspecific band in (D). The average fold increase versus the WT Rad51 level, which was set to 1.00, was calculated from 3 to 5 biological repetitions (blue numbers below the blot).

Thus, we asked whether Rad51 is modified by another ubiquitin-like modifier, SUMO. In yeast, the SUMO protein is encoded by the *SMT3* gene. Because the *SMT3* gene is essential for cell viability we could not use a null strain. First, we checked the Rad51 level in cells lacking two major SUMO E3 ligases, Siz1 and Nfi1/Siz2 (30, 31). The results showed no change in the Rad51 level in the *nfi1/siz2Δ* strain, a slight increase in the *siz1Δ* strain, and a significant increase in the double mutant *siz1Δ nfi1/siz2Δ* (Figure 3 A). Additionally, the results drew our attention to the fact that one of the bands detected specifically by anti-Rad51 antibodies (the upper Rad51 band in Figure 3 A) seemed to be a doublet in the samples derived from the *siz1Δ nfi1/siz2Δ* sample. Therefore, we applied alternative separation conditions (acrylamide:bis-acrylamide [29:1], longer gels, and slow electrophoresis), which allowed for better separation of the two upper Rad51 bands. Subsequently, in the sample derived from the *siz1Δ nfi1/siz2Δ* strain, a clear band of almost equal intensity appeared below the Rad51 band migrating around 50 kDa (Figure 3 B). Because unmodified Rad53 is predicted to migrate at approximately 43 kDa, this faster migrating Rad51 band likely corresponds to another form of post-translationally modified Rad51. The character of this modification is the subject of a separate study.

Because Rad51 accumulated in *siz1Δ nfi1/siz2Δ* cells, we asked if overproduction of SUMO E3 ligase would promote Rad51 ubiquitination. To answer this question, we overproduced the SUMO E3 ligase Siz1 in the cells, which simultaneously expressed a His-tagged ubiquitin. In particular, we used a WT strain carrying two plasmids: A plasmid under the control of the inducible *CUP1p* promoter encoding a His-tagged ubiquitin variant (24) or HA-tagged ubiquitin variant (32) (as a negative control), and a plasmid bearing the *SIZ1* gene under the *tet2Op* promoter or the empty vector pCM188 (33) as a control. Then, by using Ni^2+^-NTA sepharose, we purified ubiquitinated proteins from these cells. Western blot analysis performed using anti-Rad51 antibodies detected enrichment in poly-ubiquitinated Rad51 forms among the Ni^2+^-NTA-bound protein fraction in the sample derived from the strain overproducing Siz1 compared to the control (Figure 3C). In parallel, a decrease in Rad51 level was observed in the respective total extract sample. Together, these results led us to conclude that the Rad51 protein is poly-ubiquitinated in a Siz1- and SUMO-dependent fashion.

Several enzymes have SUMO E3 ligase activity in yeast. Besides the Siz1 and Nfi1/Siz2 SUMO E3 ligases mentioned above, a similar activity is displayed by other enzymes, such as Cst9/Zip3 or Mms21. Thus, we asked whether removing their activity would affect the Rad51 level. In the *zip3Δ* strain, we found no difference in the Rad51 level compared to the WT strain (Figure 3 D). However, in the *mms21-1* mutant (34, 35) the Rad51 level was considerably increased (Figure 3 E).

Because various SUMO E3 ligases seem to keep the Rad51 level under control, facilitating or enabling Rad51 ubiquitination, we asked whether the presence of the SUMO chain is engaged in this process. Usage of an *smt3AllR* (36) mutant, in which all lysine residues were replaced with arginines, preventing lysine-linked chain formation, enabled us to test this hypothesis. Indeed, the level of Rad51 in the *smt3AllR* mutant was significantly increased (Figure 3 E).

The SUMO chain may recruit the SUMO-targeted ubiquitin E3 ligases (STUbLs), which modify proteins to direct them for degradation. One such enzyme is the Slx5-Slx8 complex. We asked whether this complex contributes to the Rad51 level limitation. We found that the Rad51 level was raised in the *slx8Δ* mutant compared to the control (Figure 3 A). All these results confirmed our hypothesis that SUMOylation is involved in Rad51 degradation.

### Multiple enzymes regulate the ubiquitination of Rad51 in vivo

An increased Rad51 level was detected in strains lacking various E2 and E3 ligases (Figure 2). Which of them is then responsible for the ubiquitination of Rad51? To address this issue, we used two opposite approaches. First, we looked for E2 or E3 activities, whose absence resulted in the disappearance of Rad51 ubiquitinated forms from the cell. Second, we looked for enrichment of ubiquitinated forms of Rad51 when specific E2 or E3 enzymes were overproduced; we reasoned that increased activity in the conjugating and ligase enzymes would overwhelm the capacity of the degradative system, so that the ubiquitinated substrate would accumulate. In these experiments, we again took advantage of the His-tagged ubiquitin system. After introducing a YEp181-CUP1-His-Ubi plasmid (37), expression from the *CUP1p* promoter was induced with Cu^2+^ ions. Then, proteins were extracted from the cells, and the His-ubiquitin-bound protein fraction was purified using Ni^2+^-NTA sepharose and analyzed by western blot using anti-Rad51 antibodies. Ubiquitinated Rad51 forms were absent in the samples prepared from *rad6Δ* and *dia2Δ* strains (Figure 4 A), pointing to the importance of Rad6 (E2) and Dia2 (E3) for their formation.

**Figure 4.**
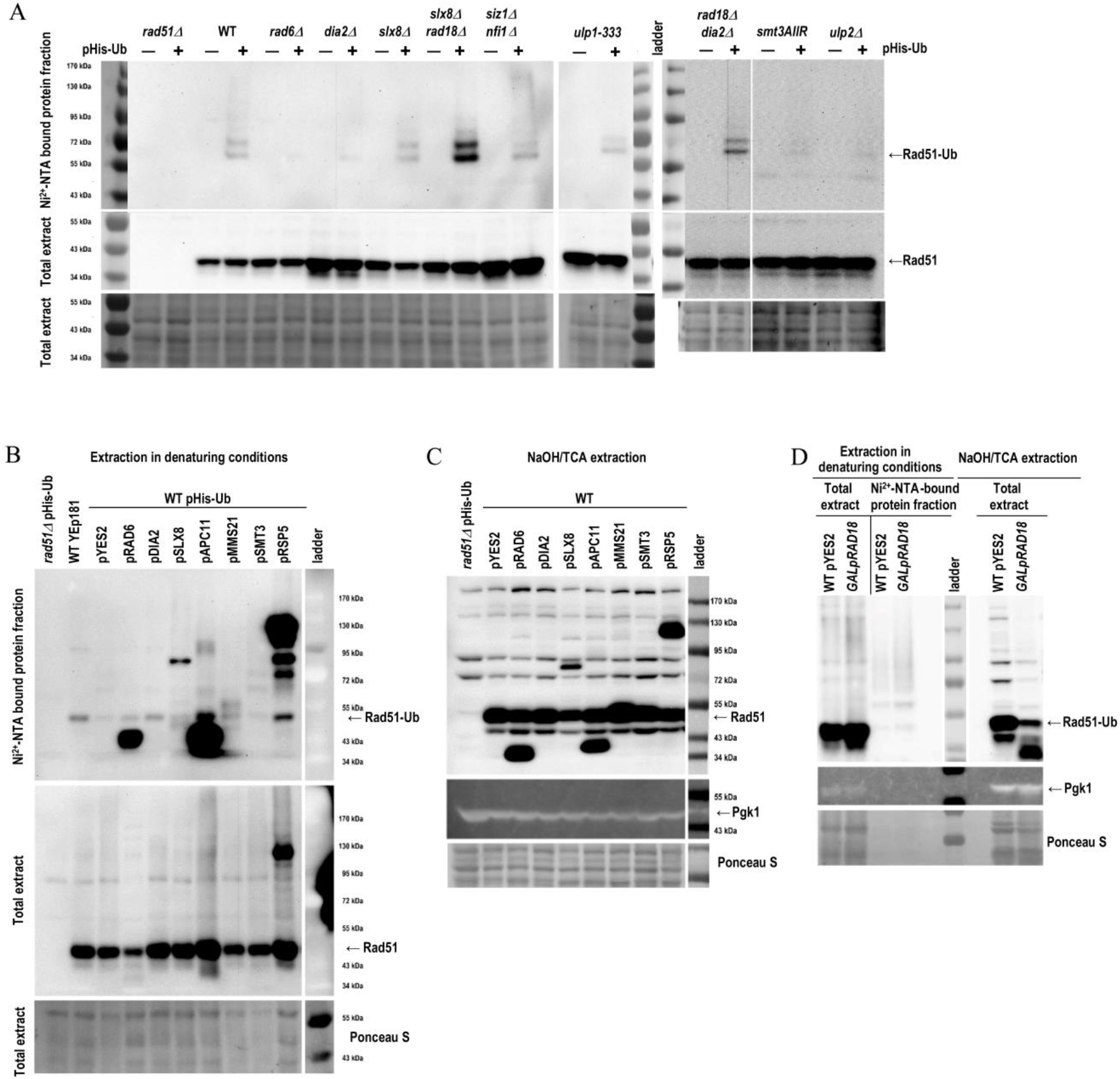
Many enzymes affect the ubiquitination pattern of Rad51. (**A**). Analysis of Rad51 ubiquitination by His-ubiquitin affinity-isolation assay using strains with defects in the ubiquitination or SUMOylation pathways. Strains in the BY4741 background bearing the YEp181-CUP1-His-Ubi^37^ plasmid were grown to exponential phase. The production of His-tagged ubiquitin was induced with Cu^2+^ ions and cells were collected. Ubiquitinated proteins were purified on a Ni^2+^-NTA Sepharose column and analyzed by western blot using anti-Rad51 antibodies. Rad51 and poly-ubiquitinated Rad51 (Rad51-Ub) bands were indicated on the right. Molecular mass markers were shown on the left. At the bottom, the blot containing total extracts stained with Ponceau S is shown. (**B**) Analysis of Rad51 ubiquitination by His-ubiquitin affinity-isolation assay using strains overproducing the proteins involved in the ubiquitination and SUMOylation pathways. Strains bearing two plasmids, enabling the production of His-tagged ubiquitin (YEp181-CUP1-His-Ubi) or pYES2 (control), or plasmids from the MORF collection as indicated in the WT (Y258) background, as well as additional control strains with the YEp181 plasmid and the *rad51Δ* strain with the YEp181-CUP1-His-Ub plasmid were grown to exponential phase. Production of His-tagged Ub and the indicated proteins was induced by Cu^2+^ ions and galactose, respectively. The ubiquitinated proteins purified on Ni^2+^-NTA sepharose were subject to SDS-PAGE, transfer, and western blot analysis with anti-Rad51 antibodies. At the bottom, the result of Ponceau S staining of a blot containing total cell extracts is shown. (**C**) Analysis of Rad51 level in the strains overproducing the proteins involved in the ubiquitination and SUMOylation pathways. The same strains as in (**B**) but without plasmid encoding His-tagged Ub were grown to exponential phase. Induction of MORF plasmid expression was performed by 4-h cultivation on galactose. Collected cells were subject to NaOH/TCA extraction, and samples were resolved on SDS-PAGE, transferred to a membrane, and subjected to western blot analysis to detect Rad51. At the bottom, the result of western blot using anti-Pgk1 antibodies and a blot containing total extracts stained with Ponceau S are shown. (**D**) Analysis of Rad51 ubiquitination by His-ubiquitin affinity-isolation assay using a *GAL1p-RAD18* (YAM27) and WT (BY4741) strain carrying plasmid YEp181-CUP1-His-Ub. The assay was performed as in (**B**).

Because we expected SUMO-targeted Rad51 ubiquitination, we included in our analysis strains defective in SUMO-chain formation (*smt3AllR*), lacking functional SUMO E3 ligases (*mms21-1, siz1Δ nfi1/siz2Δ*) or SUMO peptidases (*ulp1-333* and *ulp2Δ*). The ubiquitinated forms of Rad51 were absent or reduced substantially in the *smt3AllR* sample (Figure 4 A). Thus, the inability to form SUMO chains or lack of appropriate enzymatic activity affects Rad51 ubiquitination.

Elevated Rad51 ubiquitination might be expected in the *ulp2Δ* sample due to the lack of SUMO peptidase activity. The virtual absence of Rad51 ubiquitination in this sample may be explained in several ways: (i) SUMOylated Rad51 is a substrate for peptidases other than the Ulp2 isopeptidase; (ii) the accumulation of Rad51-SUMO forms in the *ulp2Δ* strain, stimulates a deubiquitinase, e.g., Wss1 (38), which removes ubiquitin chains from that protein; (iii) SUMO influences Rad51 ubiquitination indirectly, via some Ulp2 substrates, e.g., ubiquitin E3 ligase or SUMO E3 ligase, which in turn affect the Rad51 ubiquitination state. In the latter case, the higher rate of Rad51 ubiquitination in the strain overproducing Siz1 (Figure 3C) would be the result of activation of such an E3 ligase by SUMOylation in a Siz1-dependent fashion.

Among the analyzed strains lacking ubiquitin E3 ligases, only the *dia2Δ* strain showed a reduction in the formation of Rad51 ubiquitinated forms (Figure 4 A). A similar effect was visible in the *rad6Δ* strain lacking a ubiquitin-conjugating enzyme (E2). Deleting the genes encoding individual ubiquitin E3 ligases led to, at most, slight differences in the intensity of the higher molecular mass bands representing poly-ubiquitinated forms of Rad51 (Figure 4 A). A similar effect was seen in the double deletion *rad18Δ dia2Δ* strain; however, when we eliminated the activity of two STUbLs, Slx8 and Rad18, we found enrichment in Rad51-ubiquitin derivatives. Thus, abolishing one pathway marking Rad51 for degradation apparently activates the alternative one. We concluded that because a high level of Rad51 is toxic to the cells, backup pathways evolved to keep it under control. It is worth mentioning here that the combination *slx8Δ dia2Δ* is synthetically lethal (39). We think that at least one of the Rad51 ubiquitination pathways has to work to preserve cell viability.

In the second line of experiments, we employed the strains from the moveable ORF library (MORF collection) (40). These strains carry plasmids expressing various yeast genes from the *GAL1* promoter, which allows overproduction of selected proteins when strains grow in the presence of galactose. In our approach, we overexpressed genes of the selected E2 and E3 enzymes in the strains also producing His-ubiquitin. This approach again allowed for purification of the His-ubiquitin-tagged protein fraction on Ni^2+^-NTA sepharose and further detection of ubiquitinated Rad51. The anti-Rad51 antibody recognizes mono-ubiquitinated Rad51 in all of the analyzed strains. The pattern of poly-ubiquitination of Rad51 varies between strains; however, the control sample (WT + empty vector pYES2) and Smt3 sample (WT + pSMT3) showed a similar pattern, but the intensity of bands was increased in the latter (Figure 4B). It seems that better access to SUMO promotes Rad51 ubiquitination. The most pronounced signals of ubiquitinated Rad51 were visible in the cells overproducing Rsp5 ubiquitin E3 ligase, testifying that Rsp5 is responsible for Rad51 ubiquitination. This result, together with the reduced level of Rad51 in *rsp5-1* and *rsp5-3* mutants (Figure 2 A), raises a question concerning the role of Rsp5-dependent Rad51 ubiquitination, suggesting a role other than tagging the protein for degradation. It is worth Noting that when only Rsp5 was overproduced, but not ubiquitin, the level of the monomeric form of Rad51 (the band migrating at about 50 kDa) was lower compared to the situation when both proteins were overproduced (Figure 4 C). Note that protein extraction in denaturing conditions from the cells overproducing ubiquitin allows better preservation of the Rad51-ubiquitin derivatives. In contrast, extraction using the NaOH/TCA method from the cells expressing endogenous ubiquitin allows better visualization of protein turnover.

In the samples prepared from the cells overproducing the Slx8 subunit of the Slx5-Slx8 STUbL complex, the most intense band migrated at about 86 kDa (Figure 4 B). However, other Rad51-ubiquitin derivatives were also present in this sample; among them those visible in the sample prepared from cells expressing the gene encoding Mms21 SUMO E3 ligase. In the samples prepared from cells expressing the Rad6 ubiquitin-conjugating enzyme, the poly-ubiquitinated forms of Rad51 were hardly visible. Instead, the band migrating faster than the Rad51 monomeric band appeared, which indicated that poly-ubiquitinated Rad51 derivatives were short-lived in this strain leading to effective Rad51 degradation. Therefore, we concluded that Rad6 E2 activity contributes to Rad51-ubiquitin tagging for degradation. Also, the overproduction of Apc11, the catalytic subunit of the APC complex (ubiquitin E3 ligase), led to efficient Rad51 degradation (Figure 4 C). However, the final digestion product migrated slower than that observed for cells overproducing Rad6. Moreover, the level of the monomeric form of Rad51 in the cells overproducing Apc11 did not drop down as was seen in the case of Rad6 overproduction (Figure 4 B) but was higher than in the WT control. These results suggest the existence of different proteases that contribute to Rad51 digestion. Overall, Rad51 level control seems to be a very complex issue.

The well-known partner of Rad6 (E2) is Rad18 (E3). To get an answer to the question of whether Rad51 ubiquitination depends also on Rad18 function, we prepared a yeast strain in which a *GAL1p::RAD18* fusion (41) replaced the genomic copy of *RAD18*; *RAD18* was expressed from the genome, but its expression was dependent on the presence of galactose in the media as a sole carbon source. We also introduced the YEp181-CUP1-His-Ub plasmid to this strain allowing us to take advantage of the different methods of protein extraction and differential level of production of the substrate for the ubiquitination reaction. After 4 h of growth on galactose and in Cu^2+^ ion-containing media, the ubiquitinated proteins were purified as above. Rad51 derivatives were visualized following SDS-PAGE separation with anti-Rad51 antibodies (Figure 4 D). We found that Rad51 was ubiquitinated in a Rad18-dependent fashion (left panel). Moreover, this ubiquitination led to Rad51 degradation (right panel).

### Rad51 is SUMOylated

Because the data described above pointed to the involvement of SUMOylation in the process of Rad51 level regulation, the question arose as to whether Rad51 is SUMOylated directly or whether the observed effects were the consequences of the SUMO-dependent regulation of enzymes influencing Rad51 stability. To answer this question, we performed the analysis of the Rad51 modification using protein immunoprecipitation with anti-Rad51 antibodies and detection of SUMO and ubiquitin derivatives among immunoprecipitated forms of Rad51. We found that Rad51 was SUMOylated in the samples purified from the strain overproducing the SUMO E3 ligase Mms21 and SUMO-ligase/SUMO-targeted metalloprotease Wss1 (Figure 5, anti-Smt3). Different patterns of higher mass bands were detected in the samples derived from strains overproducing Slx8 and Rsp5 ubiquitin E3 ligases and Ulp1 protease, which cleaves specifically SUMO conjugates. In the latter sample, the lower mass form recognized by both anti-Rad51 antibody and anti-Smt3 antibody was also detected. The band of a similar mass was also seen in the sample derived from strain overproducing Apc11. In the sample derived from the strain overproducing Rad6, the band migrating even faster than that in the sample derived from strain overproducing Apc11 was visible. Interestingly, using an anti-ubiquitin antibody, we were able to show the poly-ubiquitination of Rad51. The poly-ubiquitinated forms of Rad51 were well apparent in the sample derived from strain overproducing Apc11 or Slx8; however, in the latter case, they migrated more slowly and were more pronounced (Figure 5). This experiment showed that Rad51 undergoes two posttranslational modifications, ubiquitination, and SUMOylation. Furthermore, it seems in some conditions, both modifications could be introduced simultaneously.

**Figure 5.**
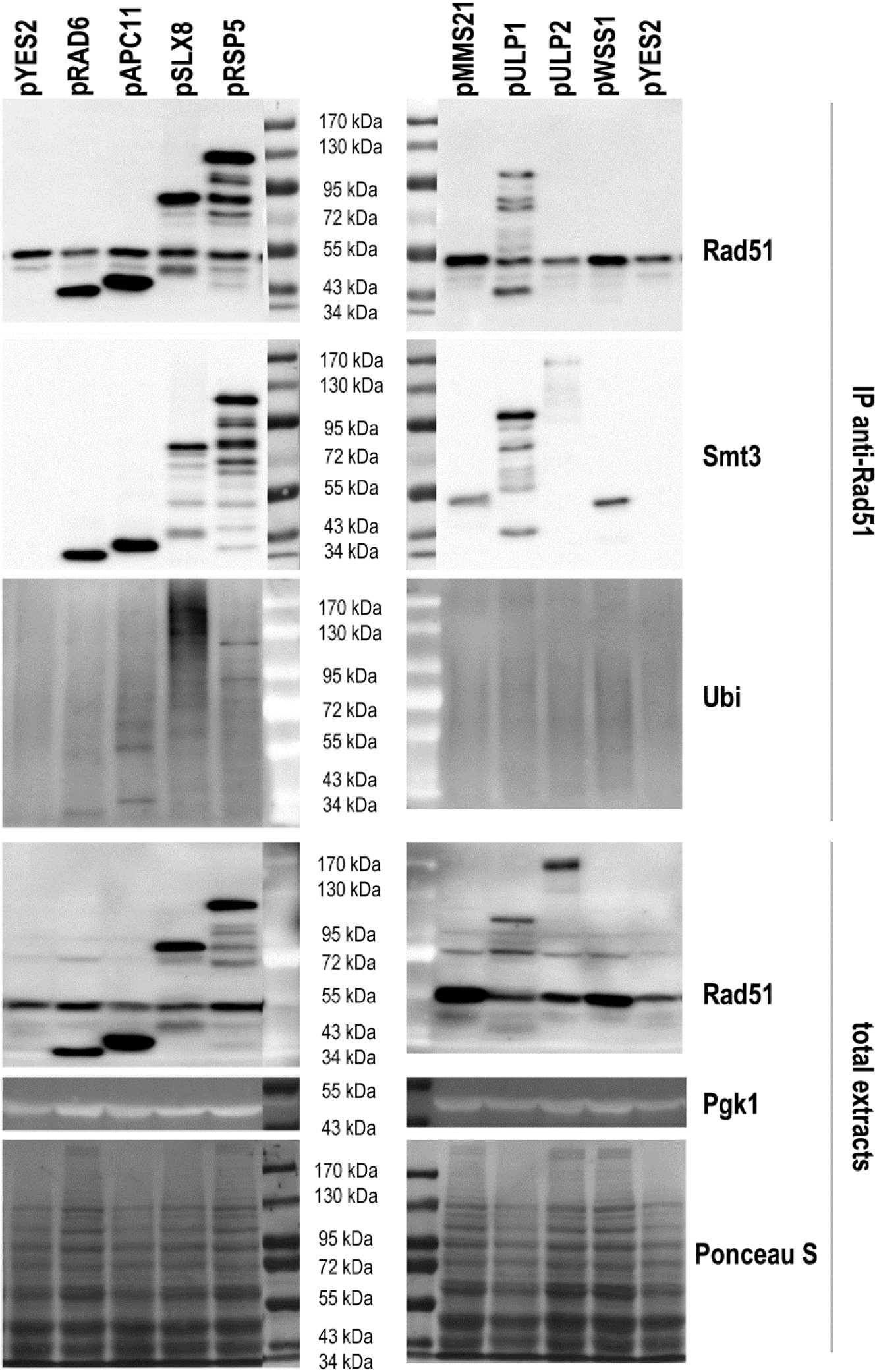
Rad51 is SUMOylated *in vivo*. Strains in the Y258 background, from the MORF collection, carrying the indicated expression plasmids or the pYES2 empty vector as a control, were cultivated to early exponential phase on SC-ura medium. Then the cells were harvested, washed, and allowed to grow for 4 h in a selective medium supplemented with 2% galactose to induce expression from the *GAL1* promoter. Protein extracts were prepared and used for Rad51 immunoprecipitation as described in the Materials and Methods section. The immunoprecipitated proteins and total extracts were subject to SDS-PAGE, transfer, and western blot with appropriate antibodies as indicated on the right side of the images. Because anti-Rad51 antibodies developed in rabbit was used for IP, we used the mouse anti-ubiquitin antibodies and rabbit antibodies conjugated with HRP for Rad51 and Smt3 protein detection. On the total extracts blot, detection of proteins of interest was performed with rabbit anti-Rad51 antibody and mouse anti-Pgk1 antibody. At the bottom, the Ponceau S-stained blot with total extracts is shown.

## Discussion

Rad51 recombinase plays such a crucial role in the cell that its level and activity have to be tightly regulated. Both depletion of Rad51 and its high level leads to a series of defects in the DNA damage response (7, 21, 42, 43). Lack of Rad51 in higher eukaryotes results in embryonic lethality or cell death (9, 44). In humans, a high level of RAD51 it is a bad prognostic in various cancers (45, 46). During DNA damage repair Rad51 acts through formation of a filament on ssDNA. Forming the nucleofilament is essential for the initial steps of homologous recombination, including DNA sequence homology search, recruitment of DNA repair proteins and protein complexes involved in chromatin remodeling, which together enable repair of DNA damage such as DNA breaks or cross-links (47).

In our work, we aimed to uncover the regulation of Rad51 level in the cell and indicate the factors contributing to this regulation. The level of Rad51 protein was increased in *pre2-1, pre3-1, pup3-1*, and *ump1Δ* mutants, suggesting proteasomal degradation of this protein. In agreement with this result, we showed Rad51 poly-ubiquitination. We found that the Rad51 level is actively regulated by proteasomal degradation, and its half-life is decreased after genotoxic stresses (Figure 1). Similar observation was made by Woo and colleagues when they used the proteasome inhibitor MG132 to determine if Rad51 is degraded by proteasome during the DNA damage response (48). We also identified several ubiquitin-conjugating enzymes (E2) and ubiquitin ligases (E3) affecting the Rad51 cellular level. The level of Rad51 is elevated in *rad6Δ* and *ubc13Δ*, as well as in *rad18Δ* and *rad5Δ* (strains lacking E2 and E3 enzymes, respectively). Besides RING finger-type E3 ubiquitin ligases (Rad5 and Rad18), also F-box proteins, Dia2 and Cdc4, subunits of SCF^Dia2^ and SCF^Cdc4^ ubiquitin ligase complexes, respectively, seems to regulate Rad51 level because, in *dia2Δ* and *cdc4-1* mutant cells, the level of Rad51 was doubled compared to the control strain level (Figure 2). Additionally, we found that the SUMO-targeted ubiquitin ligase, the Slx5-Slx8 complex, contributed to ubiquitin-dependent Rad51 regulation. In accordance with this result, we found an increased level of Rad51 protein in an *mms21-1* mutant, as well as in a *siz1Δ nfi1/siz2Δ* strain lacking two other yeast SUMO ligases. Also, the poly-ubiquitinated forms of Rad51 were enriched in cells overproducing the SUMO E3 ligase Siz1 or Smt3 (a yeast SUMO variant). Thus, Rad51 degradation is ubiquitin and SUMO dependent (Figure 3, 4, 5).

By showing the influence of E3 ubiquitin and SUMO ligases overproduction on the basal level of Rad51 and the appearance of higher molecular weight Rad51 forms in the cell, we indicated which of them are involved in regulating the Rad51 level. But we also showed that at least some of these enzymes introduce post-translational modifications to Rad51. We also showed that at least some of these enzymes introduce post-translational modifications to Rad51. The fact that Rad51 is SUMOylated and ubiquitinated *in vivo* was established. The SUMOylation of Rad51depends on Mms21, and Wss1 SUMO E3 ligases (49) (Figure 5). The ubiquitination of Rad51 depends on Rad6-Rad18, Dia2, Apc11, Slx8, and Rsp5 ubiquitin E3 ligases (Figure 4, Figure 5). However, the modification network seems to work in a complex way with respect to Rad51 stability. Modification with ubiquitin or SUMO appears to be bifunctional. In some circumstances it leads to Rad51 degradation, whereas in the others it stabilizes the protein. In addition, some of the observed effects seem to be indirect, e.g., the role of Siz1 E3 SUMO ligase in Rad51 stability. On the one hand, we showed a decreased level of Rad51 monomers and increased poly-ubiquitination of Rad51 when Siz1 was overproduced (Figure 3 C). On the other hand, we did not see the SUMOylated form of

Rad51 among immunoprecipitated Rad51 derivatives, nor among a SUMOylated protein fraction enriched on the Ni^2+^-NTA sepharose from the cells overproducing both His-tagged Smt3 and Siz1 (data not sown). We postulate that Siz1 SUMOylates the proteins that contribute to Rad51 stability. At least two such proteins could be implicated. Rsp5 is subject to Siz1-dependent SUMOylation, which results in its reduced ubiquitin ligase activity (50). Because Rsp5 stabilizes Rad51, the activity of Siz1 will stimulate Rad51 degradation. The other Siz1 substrate is Pol30, the yeast proliferating cell nuclear antigen (PCNA), the ring-shaped trimeric complex that encircles DNA and functions as a sliding clamp and processivity factor for replicative DNA polymerases (50-52). SUMOylated PCNA recruits Srs2 and Rad18, activating Rad51 translocase activity and ubiquitin ligase activity, respectively, which both act in an anti-recombinogenic manner and likely influence further Rad51 cellular fate (41, 53-56). We do not know what is happening with Rad51 protein stripped-off from DNA by the Srs2 helicase/translocase, but we showed that Rad51 might be SUMOylated and that one of the STUbLs involved in its poly-ubiquitination is Rad18 (Figure 4).

The monomeric form of Rad51 detected on western blots with the specific anti-Rad51 antibody migrates as two or three bands (dependent on the strain background), with a molecular mass of about 43 kDa (the theoretical mass of Rad51), 50 kDa (likely a monoubiquitinated Rad51), and 52 kDa (likely a monoSUMOylated Rad51). Indeed, the monoubiquitinated Rad51 was detected among the Ni^2+^-NTA-bound protein fraction of His-ubiquitin-tagged proteins in almost all analyzed strains from the MORF collection overproducing E3 ubiquitin ligases. The most pronounced signal was visible in the samples derived from strains overproducing Apc11, a catalytic subunit of the APC complex (i.e., a ubiquitin ligase, whose activity enables metaphase/anaphase transition during mitosis via tagging with ubiquitin the anaphase inhibitors leading to their degradation) and Rsp5, a NEDD4 family E3 ubiquitin ligase (involved among others in such processes as MVB sorting, endocytosis or the heat shock response) (Figure 4 B). Likely due to distinct biological processes in which these two ligases are involved, the pattern of other Rad51 bands accompanying the mono-ubiquitinated Rad51 monomer is totally different. When Apc11 was overproduced, we also saw the poly-ubiquitinated forms of Rad51 (Figure 4 B, Figure 5). Still, the most prominent Rad51 band observed for this sample migrated as 43 kDa, i.e., exactly as expected for a non-modified Rad51 protein. Only a band of that mass was observed among proteins modified with the His-tagged variant of ubiquitin. Thus, it probably represents a degradation product of Rad51, shortened by about 9 kDa. When Rsp5 was overproduced, we saw additional bands of Rad51 migrating slower than mono-ubiquitinated Rad51. These bands may reflect poly-ubiquitinated forms of Rad51; however, some of them had a mass which is a duplication or triplication of the Rad51 mass. Thus, it is also possible that the bands reflect oligomers of Rad51. In such a case, the process of Rad51 oligomerization would be stimulated by ubiquitination with the ubiquitin E3 ligase Rsp5 (Figure 4 B, Figure 5). Interestingly, the upper Rad51 bands appeared not only when we detected ubiquitinated Rad51 derivatives but also when we detected SUMOylated Rad51 forms (Figure 5). However, the pattern of SUMOylated forms only partially overlaps with the pattern of ubiquitinated forms. The results led us to conclude that part of the cellular Rad51 pool might be modified with SUMO and ubiquitin or might possess mixed SUMO and ubiquitin chains. Also, the availability of substrates for protein modification and the local concentrations of E3 ligases may determine the future fate of the protein.

Acquired data pointed to three ubiquitin E3 ligases, whose activities result in the poly-ubiquitination and subsequent degradation of Rad51. Besides Apc11, we confirmed Rad18-dependent and Slx8-dependent-poly-ubiquitination of Rad51 (Figure 4D, Figure 4 B, Figure 5). Interestingly, the Rad51 degradation product detected in the samples overproducing Rad18 migrated faster than that seen in the samples overproducing Apc11. Moreover, in the samples overproducing Rad6, the Rad51 degradation product appears, which had the same size as the one detected in the sample with Rad18 overproduction. The E2 ubiquitin-conjugating enzyme Rad6 cooperates with Rad18 to monoubiquitinate PCNA at Lys164 in response to DNA damage stress (49). The difference in the size between Rad51 degradation products appearing in the diverse conditions suggests the involvement of distinct peptidases in the degradation of this protein. Why do the additional peptidases have to be involved? First, because a high level of Rad51 is harmful for the cells, the backup systems evolved that allow the elimination of excess Rad51 when this is required. Second, the Rad51 nucleofilament might be a difficult target for degradation. Possibly, it has to be disassembled piece by piece before being sent to the proteasome for final digestion. The specialized peptidases might play an important role in this process. We presume that various enzymes tagging Rad51 protein for degradation and various peptidases could be involved in its degradation at different cell cycle phases. For example, the APC complex would be necessary for Rad51 degradation during mitosis, likely during the metaphase/anaphase transition, while the Rad6-Rad18 complex would play a similar role during the S phase of the cell cycle especially during DNA damage stress.

Among ubiquitin E3 ligases that contribute to Rad51 poly-ubiquitination, there are two enzymes belonging to the SUMO-targeted ubiquitin ligases (STUbl), Rad18 and Slx8 (41, 57). However, the patterns of Rad51-derivatives purified from cells overproducing Rad18 or Slx8 were different. While overproduction of Rad18 caused an increased poly-ubiquitination of Rad51 accompanied by the appearance of the primary/major degradation product, overproduction of Slx8 led to increased poly-ubiquitination of Rad51 and the appearance of an additional protein band of about 80 kDa (Figure 4 B, Figure 5). Puzzling is the fact that an 80-kDa protein band was visible in samples from cells overproducing Slx8 both after Ni^2+^-NTA sepharose enrichment of ubiquitinated proteins and after anti-Rad51 immunoprecipitation. This band was detected both with anti-Smt3 and anti-Rad51 antibodies. Thus, this protein likely possesses both ubiquitin and SUMO modifications. Notably, the SUMO signal was more pronounced. Interestingly, polyubiquitinated forms of Rad51 released from the cells overproducing Slx8 had a mass over 80 kDa. We did not detect the degradation product of Rad51 in these cells; however, the level of Rad51 monomers was decreased in total extracts from the cells overproducing Slx8, at least when His-tagged ubiquitin was not overproduced in parallel. A protein of similar mass was also observed when Rsp5 ligase was overproduced; but in this case it was accompanied by additional Rad51 derivatives of higher mass. We wonder, what exactly is the derivative of Rad51 that has a mass higher than 80 kDa? It could it be the Rad51 monomer modified with a SUMO/ubiquitin chain or the Rad51 dimer, the formation of which was stimulated by posttranslational modification. We cannot exclude any of these possibilities yet.

This puzzle could be resolved by experiments made with the usage of *RAD51* alleles containing point mutations that prevent modifications. The problem is that there are 17 lysine and 8 cysteine residues in the Rad51 protein, which can be modified by ubiquitin or SUMO. Two reports concerning possible ubiquitination residues in Rad51 exist; both came from proteomic approaches (58,59). In both attempts, the lysine 131 (K131) residue was indicated as a ubiquitination site. Moreover, Back and colleagues showed that the K131 might also be modified by K63-linked poly-ubiquitin chains in response to oxidative stress (59). Among various types of ubiquitin chains, which might be conjugated to the protein substrates, the K63-linked poly-ubiquitin chain is thought to be the only linkage that is not necessarily related to protein degradation by the proteasome. Instead, K63-linked chains are mostly used as signals in DNA repair, trafficking and autophagy (60). Thus, the marking of Rad51 K131 with K63-linked poly-ubiquitin chains would be also connected with the modulation of protein function during genotoxic stress or cellular trafficking.

SUMOylation and ubiquitination are involved in regulating processes that are crucial for genome maintenance, such as replication, DNA repair, and chromosome segregation. However, we do not know yet how SUMOylation and ubiquitination influence the molecular activity of Rad51. Nevertheless, the data presented here permitted an initial look at this regulation circuit, which is summarized in Figure 6. The Rad51 level, and likely also its activity, is regulated by a network of ubiquitin and SUMO ligases. It is highly probable that each of these enzymes acts in certain circumstances. According to current knowledge concerning other substrates of these enzymes, we can anticipate the particular moments in the life of the cell when they play a major role in Rad51 regulation.

**Figure 6.**
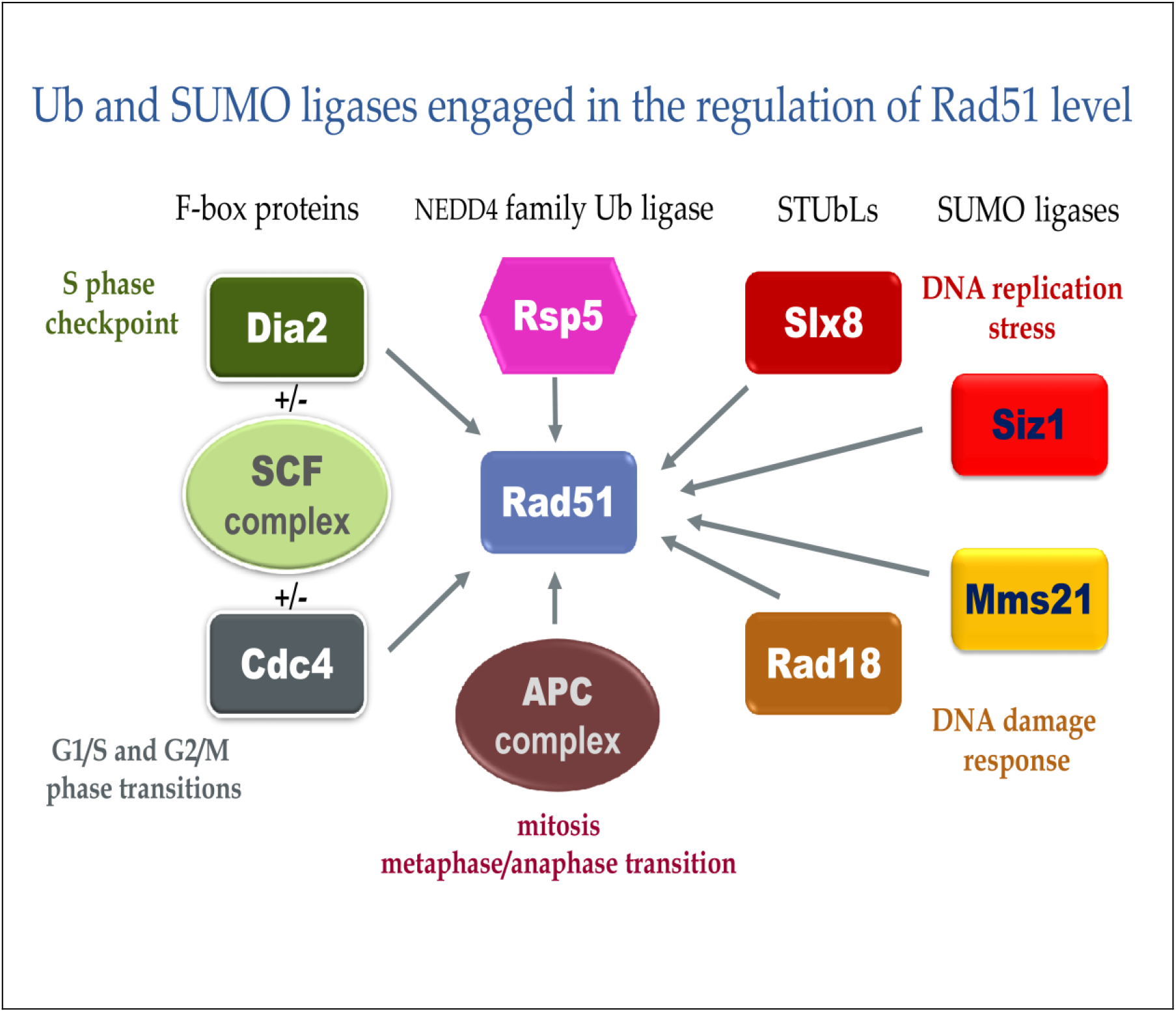
Factors that contribute to Rad51 level regulation. Ubiquitin and SUMO ligases engaged in posttranslational modification of Rad51 recombinase, influencing Rad51 level and activity. The scheme is based on published data concerning other substrates of the presented enzymes. The moments in the cellular life when a certain enzyme modifies Rad51 are indicated.

## Materials and Methods

### Strains and plasmids

Most of the strains used in this study (Table S1) are in the BY4741 (*Mat a his3Δ1 leu2Δ0 met15Δ0 ura3Δ0*) and WCG4a (*MAT*a *ura3 leu2-3,112 his3-11,15 rad5-535 Can*^*S*^ *GAL2*) background. The deletion constructs in the BY4741 background came from the yeast knockout collection (Open Biosystems), and were initially prepared during the *Saccharomyces* genome deletion project (61). Strains carrying point mutations affecting essential genes in the BY4741 background came from a yeast temperature-sensitive mutant collection (62). Strain YAD11 was constructed by gene replacement of the *SIZ1* gene by a *siz1::hphMX6* cassette, amplified using siz1up and siz1lw primers (Table S2) and genomic DNA isolated from the YAH144 (63) strain as a template, in the BY4741 *nfi1/siz2Δ* strain, using a standard *in vivo* recombination method.

Strains YAM7 and YAM8 were created by introduction of the *rad18::LEU2* and *rad18::URA3* cassettes to the BY4741 *slx8Δ* strain, respectively. The *rad18::LEU2* cassette was obtained from a PCR reaction using RAD18up and RAD18lw primers and YAS33 (64) strain genomic DNA as template. The *rad18::URA3* cassette was obtained in a PCR reaction using RAD18kup and RAD18klw primers and pRS316 (65) DNA as template. Strain YAM19 was created by introduction of the *rad18::URA3* cassettes to the BY4741 *dia2Δ* strain. Strain YAM9 was prepared by replacement of the *ULP2* gene by a *ulp2::natNT2* cassette, amplified using ULP2kup and ULP2klw primers (Table S2) and pRS41N plasmid DNA (66) (Table S3) as a template, in the BY4741 strain, using a standard *in vivo* recombination method. Strains YJK1 and YJK2 are WCG4a and YWH24 derivatives, respectively, in which the *TRP1* gene was replaced with *trp1::kanMX4* by marker swap using pM3925 (67) vector. Strain YAS27 is a BY4741 derivative obtained by introducing to the genome the *GAL1pRAD18* fusion in the *RAD18* locus. For this purpose, the YIp211-GALpRAD18 (41) plasmid digested with SalI was used. The correctness of the strain was proved by PCR using GAL1A and RAD18lw primers.

Plasmid pCM188-SIZ1 was made by PCR amplification of the DNA fragment carrying the *SIZ1* ORF with SIZpup and SIZ1n-lw primers and BY4741 genomic DNA as a template and cloning into the pCM188 (33) vector digested with PmeI.

**Table S1.** Strains used in this study.

| Name | Relevant genotype | Source |
| --- | --- | --- |
| YAH144 | *MATa ade2-1(och) his3-11,15 leu2 EcoRI::URA3::leu2-BstEI trp1-1(am) ura3-1 can1-100 RAD5<sup>+</sup> siz1::hphMX4* | (63) |
| **WCG4a derivatives** |  |  |
| WCG4a | *MATa ura3 leu2-3,112 his3-11,15 rad5-535 Can<sup>S</sup> GAL2* |  |
| YJM56 | *mms2::kanMX4* | (63) |
| YWH24 | *pre2-K108R* | (22) |
| YUS1 | *pre3-T20A* | (22) |
| YUS4 | *pup1-T30A* | (22) |
| YAS33 | *rad18::LEU2 ump::kanMX4* | (68) |
| YJM26 | *rad5::HIS3* | (63) |
| YAZ10 | *rad6::LEU2* | lab stock |
| YJK1 | *trp1::kanMX4* | This study |
| YJK2 | *trp1::kanMX4 pre2-K108R* | This study |
| YJD1 | *ufo1::LEU2* | lab stock |
| YAS13 | *ump1::kanMX4* | (69) |
| **BY4741 derivatives** |  |  |
| BY4741 | *Mat a his3Δ1 leu2Δ0 met15Δ0 ura3Δ0* | EUROSCARF |
| *atg8Δ* | *atg8::kanMX4* | Open Biosystems |
| *atg12Δ* | *atg12::kanMX4* | Open Biosystems |
| *bre1Δ* | *bre1::kanMX4* | Open Biosystems |
| *der1Δ* | *der1::kanMX4* | Open Biosystems |
| *dfm1Δ* | *dfm1::kanMX4* | Open Biosystems |
| *dia2*Δ | *dia2::kanMX4* | Open Biosystems |
| *hub1*Δ | *hub1::kanMX4* | Open Biosystems |
| *mms2*Δ | *mms2::kanMX4* | Open Biosystems |
| *rad5*Δ | *rad5::kanMX4* | Open Biosystems |
| *rad51*Δ | *rad51::kanMX4* | EUROSCARF |
| *rad6*Δ | *rad6::kanMX4* | Open Biosystems |
| *rad18*Δ | *rad18::kanMX4* | Open Biosystems |
| *rri1*Δ | *rri1::kanMX4* | Open Biosystems |
| *rtt101*Δ | *rtt101::kanMX4* | Open Biosystems |
| *rub1*Δ | *rub1::kanMX4* | Open Biosystems |
| *siz1*Δ | *siz1Δ::kanMX4* | Open Biosystems |
| *siz2*Δ | *nfi1/siz2Δ::kanMX4* | Open Biosystems |
| *slx5*Δ | *slx5::kanMX4* | Open Biosystems |
| *slx8*Δ | *slx8::kanMX4* | Open Biosystems |
| *ubc13*Δ | *ubc13::kanMX4* | Open Biosystems |
| *ubp14*Δ | *ubp14::kanMX4* | Open Biosystems |
| *ump1*Δ | *ump1::kanMX4* | EUROSCARF |
| *urm1*Δ | *urm1::kanMX4* | Open Biosystems |
| *wss1*Δ | *wss1::kanMX4* | Open Biosystems |
| *yuh1*Δ | *yuh1::kanMX4* | Open Biosystems |
| *zip1*Δ | *zip1::kanMX4* | Open Biosystems |
| YAD11 | *siz1Δ:hphMX6 nfi1/siz2Δ::kanMX4* | This study |
| YJAM9 | *ulp2::natNT2* | This study |
| YJAM7 | *slx8::kanMX4 rad18::LEU2* | This study |
| YJAM8 | *slx8::kanMX4 rad18::URA3* | This study |
| YJAM19 | *dia2::kanMX4 rad18::URA3* | This study |
| *apc11-13* | *apc11-13::kanMX4* | (62) |
| *apc11-22* | *apc11-22::kanMX4* | (62) |
| *apc2-8* | *apc2-8::kanMX4* | (62) |
| *cdc23-1* | *cdc23-1::kanMX4* | (62) |
| *cdc23-4* | *cdc23-4::kanMX4* | (62) |
| *cdc34-2* | *cdc34-2::kanMX4* | (62) |
| *cdc4-1* | *cdc4-1::kanMX4* | (62) |
| *cdc4-2* | *cdc4-2::kanMX4* | (62) |
| *cdc4-3* | *cdc4-3::kanMX4* | (62) |
| *cdc53-1* | *cdc53-1::kanMX4* | (62) |
| *mms21-1* | *mms21-1::kanMX4* | (62) |
| *pre2-1* | *pre2-1::kanMX4* | (62) |
| *pre2-2* | *pre2-2::kanMX4* | (62) |
| *pre2-75* | *pre2-75::kanMX4* | (62) |
| *prp19-1* | *prp19-1::kanMX4* | (62) |
| *rpn11-8* | *rpn11-8::kanMX4* | (62) |
| *rpn11-14* | *rpn11-14::kanMX4* | (62) |
| *rpn1-821* | *rpn1-821::kanMX4* | (62) |
| *rpt1-1* | *rpt1-1::kanMX4* | (62) |
| *rsp5-1* | *rsp5-1::kanMX4* | (62) |
| *rsp5-3* | *rsp5-3::kanMX4* | (62) |
| *ulp1-333* | *ulp1-333::kanMX4* | (62) |
| YAM27 | *GALpRAD18* | This study |
| Y7092 | *MAT alpha can1Δ::STE2pr-Sp_his5* | (36) |
| *smt3AIIIR* | *MAT alpha can1Δ::STE2pr-Sp_his5 lyp1Δ smt3AIIIR::Nat* | (36) |
| **Y258 derivatives** |  |  |
| Y258 | *Mat a, pep4-3, his4-580, ura3-52, leu2-3, 112* | (39) |
| pYES2 | *Mat a, pep4-3, his4-580, ura3-52, leu2-3, 112* pYES2 | This study |
| pAPC11 | *Mat a, pep4-3, his4-580, ura3-52, leu2-3, 112* pAPC11 | (39) |
| pDIA2 | *Mat a, pep4-3, his4-580, ura3-52, leu2-3, 112* pDIA2 | (39) |
| pMMS21 | *Mat a, pep4-3, his4-580, ura3-52, leu2-3, 112* pMMS21 | (39) |
| pRAD6 | *Mat a, pep4-3, his4-580, ura3-52, leu2-3, 112* pRAD6 | (39) |
| pRSP5 | *Mat a, pep4-3, his4-580, ura3-52, leu2-3, 112* pRSP5 | (39) |
| pSLX8 | *Mat a, pep4-3, his4-580, ura3-52, leu2-3, 112* pSLX8 | (39) |
| pULP1 | *Mat a, pep4-3, his4-580, ura3-52, leu2-3, 112* pULP1 | (39) |
| pULP2 | *Mat a, pep4-3, his4-580, ura3-52, leu2-3, 112* pULP2 | (39) |
| pWSS1 | *Mat a, pep4-3, his4-580, ura3-52, leu2-3, 112* pWSS1 | (39) |

**Table S2.**
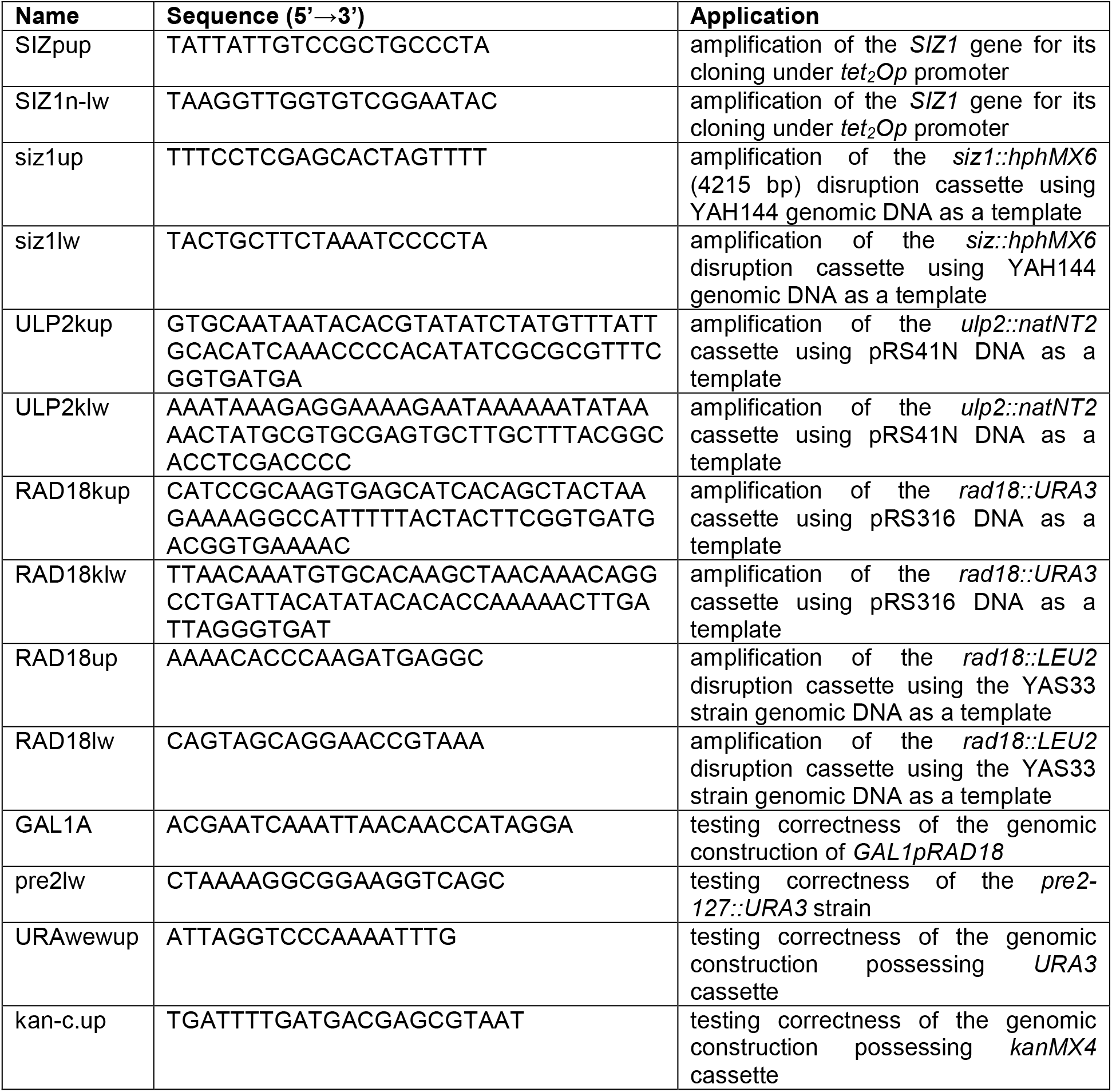
Primers used in this study.

**Table S3.** Plasmids used in this study.

| Name | Genotype | Source |
| --- | --- | --- |
| YEp96-6His-Ub | *2μ, TRP1, CUP1p-6His-UBI* | (24) |
| YEp112 | *2μ, TRP1, CUP1p-HA-UBI* | (32) |
| YEp96 | *2μ, TRP1* | (32) |
| pCM188 | *ARS-CEN, URA3, tet2Op* | (33) |
| pCM188-SIZ1 | *ARS-CEN, URA3, tet2Op-SIZ1* | This study |
| pM3925 | *trp1::kanMX4* | (67) |
| pM4758 | *kanMX4::URA3* | (67) |
| pRS41N | *ARS-CEN, natNT2* | (66) |
| pRS316 | *ARS-CEN, URA3* | (65) |
| YEp181-CUP1-His-Smt3 | *2μ, LEU2, CUP1p-His-SMT3* | (70) |
| YEp181-CUP1-His-Ubi | *2μ, LEU2, CUP1p-His-UBI* | (37) |
| YEp181 | *2μ, LEU2* | (37) |
| Ylp211-GAL-RAD18 | *URA3, GALpRAD18* | (41) |
| pYES2 | *ARS-CEN, URA3* | lab stock |

### Culture media and growth conditions

YPD medium contained 1% yeast extract (Difco, Mt Pritchard, NSW, Australia), 2% bactopeptone (Difco) and 2% glucose (POCh, Gliwice, Poland). YPGal was made as YPD, only galactose was used instead of glucose. SC medium contained 0.67% yeast nitrogen base (Difco) and 2% glucose and was supplemented with all amino acids, uracil and adenine (Formedium, Hunstanton, UK). The solid medium also contained 2.5% agar (Difco). Liquid cultures were grown with agitation at ∼200 r.p.m. (New Brunswick Scientific, Edison, NJ); the temperature sensitive mutants were grown at 23°C, and other strains at 28°C.

### Determination of protein half-life

Yeast cells were grown at 28°C in YPD medium to a density of 5 × 10^7^ cells/ml. Then, the culture was divided into three parts. The first part was treated with 0.03% MMS (Merck), the second with zeocin (Invivogen; 50 mg/ml final concentration), and the third part was the non-treated control. After 90 min of incubation at 28°C cycloheximide (Sigma, St. Louis, MO, USA; 0.5 mg*/*ml final concentration) was added. Protein samples were harvested at different time-points after treatment. Then, the level of protein in the sample was determined by western blot. Three independent biological repetitions of this experiment were performed.

### Protein analysis

For western blotting, cells were grown to the exponential phase (5×10^6^ cells/ml) in YPD medium at appropriate temperature with shaking. Then, 1×10^8^ cells were collected by centrifugation and proteins were extracted using the NaOH/TCA method. Samples were suspended in Laemmli sample buffer supplemented with a protease and phosphatase inhibitor cocktail (PIC; 1 mM PMSF, cOmplete™ Protease Inhibitor Cocktail [Roche, Base, Switzerland], PhosSTOP [Roche, Base, Switzerland]), and 10 mM N-ethyl maleimide (Sigma, St. Louis, MO, USA). Samples were vigorously vortexed and frozen in liquid nitrogen. Immediately before use, the samples were boiled for 5 min. After centrifugation (19,300 g for 2 min), equal volumes of the cell extracts were separated by SDS-PAGE (7% or 8% polyacrylamide gel, depending on the mass of the analyzed protein), and the proteins transferred onto PVDF membrane (Amersham, Germany). Blots were blocked for 2 h in 5% (w:v) nonfat dried milk and/or 3% (w:v) BSA in TBST (25 mM Tris-HCl, pH 7.5, 137 mM NaCl, 27 mM KCl, 0.1% [v:v] Tween-20) before probing with primary antibodies. Rad51 protein was detected by incubating the membranes with rabbit polyclonal antibody anti-Rad51 (1:2000, Thermo Fisher Scientific Cat# PA5-34905, RRID:AB_2552256) followed by incubation with goat anti-rabbit IgG conjugated to horseradish peroxidase (HRP) (1:2000, Agilent Cat# P0448, RRID:AB_2617138). Ubiquitin was detected using mouse monoclonal anti-ubiquitin antibody (1:500, (Santa Cruz Biotechnology Cat# sc-8017, RRID:AB_628423), yeast SUMO was detected with rabbit polyclonal anti-Smt3 antibody (1:1000, Abcam Cat# ab14405, RRID:AB_301186), followed by incubation with goat anti-mouse IgG conjugated to HRP (1:2000, Agilent Cat# P0447, RRID:AB_2617137) or goat anti-rabbit IgG conjugated to HRP (1:2000, Agilent Cat# P0448, RRID:AB_2617138), respectively. For normalization of signals, we used actin or Pgk1, which was detected by using mouse anti-actin monoclonal antibody (1:5000, (Millipore Cat# MAB1501, RRID:AB_2223041)) or mouse anti-Pgk1 antibody (1:10000, Abcam Cat# ab113687, RRID:AB_10861977), respectively, followed by goat anti-mouse IgG (H+L) alkaline phosphatase (AP)-conjugated (1:3000, Bio-Rad Cat# 170-6520, RRID:AB_11125348) antibody or Alexa Fluor 546 goat anti-mouse IgG (H+L) (1: 3000, Thermo Fisher Scientific Cat# A-11030, RRID:AB_2534089). Immunoreactive proteins on the blots were visualized using chemiluminescent substrates: SuperSignal WestPico (Pierce) for HRP and CDP Star, ready-to-use (Roche, Base, Switzerland) for AP and documented with a charge-coupled device camera (FluorChem Q Multi Image III, Alpha Innotech, San Leandro, CA). The results from independent experiments were averaged to determine the relative protein levels. The resulting bands were quantified by using Image Quant 5.2 (Molecular Dynamics, Inc., Sunnyvale, CA). The protein level was normalized to that of Act1, Pgk1 or to total proteins detected with Ponceau S staining.

### Affinity-isolation assay

The affinity isolation of His6-tagged ubiquitinated and His6-tagged SUMOylated Rad51 was performed in two biological replicates, as described previously (71). The strains in the WCG4a background (YJK1 and YJK2) were transformed with the plasmids YEp96-6His-Ubi (22), YEp112 [*UBI-HA*] or YEp96 empty vector (32). The strains in the BY4741 background were transformed with the plasmids YEp181-CUP1-His-Smt3 (70), YEp181-CUP1-His-Ubi or YEp181 empty vector (37). Transformants were grown to exponential phase (5 × 10^6^ cells/ml) at 28°C with shaking in appropriate synthetic selective media. Then, CuSO4 was added to a final concentration of 100 µM and cells were cultivated for an additional 4 h. The same number of cells (10^9^) from each culture was harvested and disrupted with glass beads in a lysis buffer (100 mM NaPi, pH 8, 10 mM Tris, pH 8, 6 M guanidine, 5 mM imidazole, 10 mM β-mercaptoethanol, 0.1% Triton X-100). The lysate was incubated with Ni^2+^-NTA beads (Ni-NTA Superflow, Qiagen) for 2 h at room temperature with rocking, and washed first with lysis buffer and, then with washing buffer (100 mM NaPi, pH 6.4, 10 mM Tris, pH 6.4, 8 M urea, 10 mM mercaptoethanol, 0.1% Triton X-100). The fraction of proteins bound to the Ni^2+^-NTA beads was eluted with a Laemmli sample buffer containing 6 M urea. All buffers, except the latter were supplemented with cOmplete protease inhibitor (Roche), PhosSTOP (Roche) and N-ethylmaleimide (NEM; SIGMA). The total lysate and Ni^2+^-NTA resin-bound proteins and non-bound supernatants were separated by SDS-PAGE and analyzed by western blot with anti-Rad51 antibody.

### Immunoprecipitation

Yeast strains from the MORF collection carrying plasmid of interest were grown on selective medium SC-URA+2% glucose at an appropriate temperature with shaking to the exponential phase (5×10^6^ cells/ml). Then cells were collected by centrifugation at 1600 g for 3 min, washed with YNB medium, suspended in SC-URA+2% galactose, and allowed to grow for an additional 4 h to induce the *GAL1* promoter. Next, 1×10^9^ cells were collected by centrifugation at 1600 g for 3 min, and proteins were extracted in 700 µl of RIPA buffer (150 mM NaCl, 50 mM Tris, pH 7.4, 1% Nonidet P-40, 0.5% sodium deoxycholate, 0.1% SDS supplemented with PIC and 1 mM Na3VO4) using the glass beads method. After centrifugation at 16000 g for 10 min at 4°C, the protein extract was transferred to a new tube. A portion of the extract was mixed with a 4 x Laemmle sample buffer (0.2 M Tris, pH 6.8, 8% SDS, 20% β-mercaptoethanol, 40% glycerol, 0.02% [w:v] bromophenol blue) for control of protein levels in the total extract. The immunoprecipitation was performed for 1 h at 4°C with rotation using 500 μl of protein extract, 300 μl of WASH buffer (150 mM NaCl, 50 mM Tris, pH 7.5 supplemented with PIC and 1 mM Na3VO4) and 50 μl of rabbit anti-Rad51 antibody (Thermo Fisher Scientific Cat# PA5-34905, RRID:AB_2552256) covalently linked to the magnetic agarose resin (CNBr-activated StepFast MAG, Bio Toolomics Ltd. Consett, County Durham, UK) according to the supplier’s protocol. After incubation, the resins were washed three times with 1 ml of WASH buffer using a magnetic rack for resin separation. Finally, the immunoprecipitated proteins were suspended in 90 μl of 1.5 x Laemmli sample buffer supplemented with PIC and 1 mM Na3VO4). Samples were frozen in liquid nitrogen and stored at -70°C until further usage. Prior to SDS-PAGE, samples were heated at 99°C for 5 min in a heatblock. After SDS-PAGE, wet transfer in Towbin buffer (25 mM Tris base, 192 mM glycine) was performed overnight at 4°C on a PVDF membrane (Amersham). The western hybridization with appropriate antibodies was then performed.

## Acknowledgments

We thank Brenda J. Adrews, Ryszard Korona and Ewa Sledziewska-Gójska for strains, and Helle D. Ulrich and Teresa Zoladek for plasmids.

## Financing

This work was supported by the National Science Center grant 2016/21/B/NZ3/03641 and Institute of Biochemistry and Biophysics, Polish Academy of Science grant DEC-MG-1/21-16 to A.S.

